# Autocrine signaling explains the emergence of Allee effects in cancer cell populations

**DOI:** 10.1101/2021.07.14.452320

**Authors:** Philip Gerlee, Philipp M. Altrock, Cecilia Krona, Sven Nelander

## Abstract

In many human cancers, the rate of cell growth depends crucially on the size of the tumour cell population. Low, zero, or negative growth at low population densities is known as the Allee effect; this effect has been studied extensively in ecology, but so far lacks a good explanation in the cancer setting. Here, we formulate and analyze an individual-based model of cancer, in which cell division rates are increased by the local concentration of an autocrine growth factor produced by the cancer cells themselves. We show, analytically and by simulation, that autocrine signaling suffices to cause both strong and weak Allee effects. Whether low cell densities lead to negative (strong effect) or reduced (weak effect) growth rate depends directly on the ratio of cell death to proliferation, and indirectly on cellular dispersal. Our model is consistent with experimental observations of brain tumor cells grown at different densities. We propose that further studying and quantifying population-wide feedback, impacting cell growth, will be central for advancing our understanding of cancer dynamics and treatment, potentially exploiting Allee effects for therapy.

## INTRODUCTION

Cancer growth is increasingly understood as an ecosystem, in which the different cellular components not only grow, but also interact. Such cellular interactions were originally proposed by Laird [1] and the subsequent discovery of growth factors provided a mechanism by which the interactions can occur [2–5]. In the 1970s, Bronk et al [6] showed that dependence on growth factors can give rise to a latent period preceding exponential growth. This suggested that negative feedback at low population density can interfere with exponential tumor expansion. Until recently, however, comparatively little attention has been given to developing mathematical models for these phenomena. The lack of modeling effort may partly reflect the high complexity of cancer cell populations [7], which makes it hard to quantify to what degree functional interactions among cells cause deviations from overall exponential growth [8, 9]. Yet, recent methodological advances make it possible to quantify how interactions between distinct subclones within cell populations affect the growth dynamics of the tumor as a whole [10, 11]. These observations motivate the formulation of mathematical models for how growing cancer cells deviate from exponential growth in a non-linear manner.

One such nonlinear growth behavior, with strong empirical support, is the Allee effect. A central idea in ecological population dynamics, the Allee effect denotes a per-capita growth rate that is reduced at low population densities [12]. There is a distinction between a weak Allee effect, for which the per-capita growth rate increases but remains positive for all densities, and a strong Allee effect where the growth rate becomes negative for sufficiently low densities before approaching zero. In the latter case, there exists a critical population density below which the population will likely go extinct. Therefore, the strong Allee effect has been studied extensively in the context of ecology and species conservation [13]. In theoretical ecology, proposed mechanisms for this include mate limitation (the problem of finding a mate at low population densities), cooperative defense, and cooperative feeding [14–17].

Recently an Allee effect was observed in cancer cell populations cultured in *in vitro* conditions in the lab at limiting densities [18]. Also, strong Allee effects are suggested by *in vivo* xenograft mouse models, where the number of xenotransplanted cancer cells need to exceed a threshold density for a tumor (or metastasis) to form [19]. This threshold depends on the type of tumor cell injected, the host animal strain and site of injection. The possibility of directly observing an Allee effect in human tumors is limited, since the effect is only present at population densities well below the clinical detection threshold at which the tumor typically contains on the order of 10^9^ cells [20]. However, by considering the rate and timing of recurrence after surgery it has been suggested that a weak Allee effect is present among cancer cells that form glioblastomas, a particular form of brain tumor in adults [21]. This conclusion was supported by a mathematical model that describes the tumor growth post-resection, in which the recurrence after surgery were better explained by assuming a weak Allee effect. Further, their results showed that cultured glioblastoma cells indeed exhibited a weak Allee effect.

Autocrine growth factor signaling could be a likely mechanism behind Allee effects in cancer cell populations. Diffusive signaling molecules released by the cancer cells themselves subsequently bind to cell surface receptors, which triggers a signaling cascade ultimately leading to the up-regulation of cell division. In glioblastoma one such growth factor is platelet-derived growth factor (PDGF), which is known to be regularly produced and to up-regulate cell division among glioblastoma cells [22]. This mechanism leads to a type of cooperative behavior and should intuitively lead to an Allee effect.

Here we show for the the first time that autocrine signaling in an *in vitro* system can give rise to an Allee effect. We study a hybrid individual-based (IB) model that describes the cells as discrete entities and the secreted autocrine growth factor as a continuous field. Further, using analytical tools [23] we derive a mean-field ordinary differential (ODE) model for the cell density, which exhibits both a weak and strong Allee effect depending on the ratio of the rates of cell birth to cell death. Lastly, we fit the ODE-model to *in vitro* growth data of glioblastoma cell culture growth and show that an Allee effect is present.

## METHODS

### Individual-based model

In order to model the effects of autocrine signaling we consider an individual-based (IB)-model in which the cells reside on a two-dimensional square lattice (see fig. 1 (a) and [23] for details). The linear size of the domain is *L* = 1 cm and it thus contains *N* × *N* cells, each with a diameter *d* = *L*/*N*. For cancer cells a typical value is *N* = 200, which gives a cell size of *d* = 50 *μ*m [24]. The growth factor (GF) concentration evolves according to

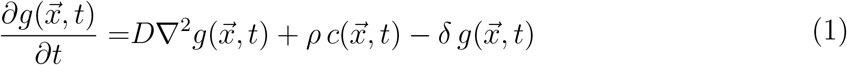

where 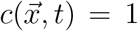 at all sites that are occupied by cells and zero otherwise. The growth factor diffuses with diffusion constant *D*, is produced at rate *ρ* and decays at rate *δ*. The partial differential equation is subject to no-flux boundary conditions, representing a closed experimental system.

**FIG. 1.**
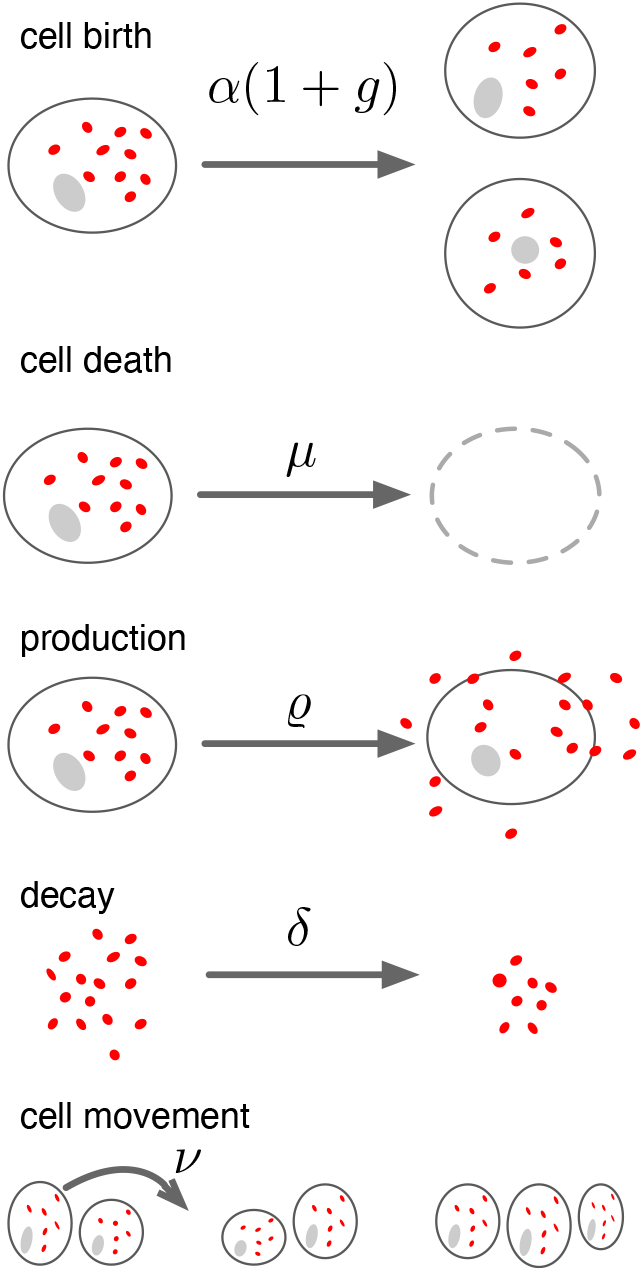
Overview of the mechanisms assumed in the model. Cancer cells divide at a rate *α*(1 + *g*), where *g* is the local growth factor (GF) concentration, and die at a constant rate *μ*. The GF is produced at rate *ρ* by all cancer cells and decays at rate *δ*. Lastly, cells migrate at rate *ν*. Parameter values are provided in Table I.

**TABLE I.** Model parameters

|  |  |  |  |
| --- | --- | --- | --- |
| $\alpha$ | Baseline cell division rate | $10^{-5}$ [25] | $\text{s}^{-1}$ |
| $\mu$ | Rate of cell death | $10^{-6} - 10^{-4}$ [26] | $\text{s}^{-1}$ |
| $\delta$ | Growth factor decay rate | $10^{-2}$ [27] | $\text{s}^{-1}$ |
| $\rho$ | Growth factor production rate | $10^{-1}$ [28] | $\text{s}^{-1}$ |
| $D$ | Growth factor diffusion coefficient | $10^{-6}$ [29] | $\text{cm}^2/\text{s}$ |
| $\nu$ | Cellular jump rate | $4 \times 10^{-4}$ [30] | $\text{s}^{-1}$ |
| $L$ | Size of the domain | 1 | cm |
| $d$ | Cellular diameter | 50 [24] | $\mu\text{m}$ |

The cell population changes in the following way: A cell located at site 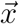 divides at a rate

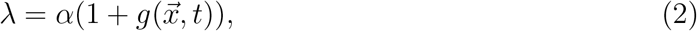

where 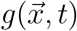 is the local GF-concentration. We consider two different modes of dispersal upon cell division. For long range dispersal the daughter is placed uniformly at random among all sites on the lattice, whereas for short range dispersal the daughter is placed uniformly at random among neighbouring lattice sites. In both cases, if the chosen site is occupied cell division fails. Cells are assumed to die at a constant rate *μ*, and also move into empty neighboring sites at rate *ν*. Movement occurs at random into neighboring sites and fails if the target site is occupied. An overview of the model is shown in fig. 1. All parameters are given in Table I.

### Analytical results

In order to understand the dynamics of the IB-model we employ a technique developed by Gerlee & Altrock [23], which makes it possible to derive an ordinary differential equation for the density of cells. This derivation is only exact in the case of long range dispersal, but we will also compare the analytical results with dynamics of short range dispersal. The main idea behind the method is to represent the stochastic distribution of cells in the IB-model as a Fourier series and from that compute the expected growth rate. The model in that paper described a situation where the population consists of two distinct subpopulations: producers that produce a diffusible public good that is costly to produce and free-riders that are identical to producers except they do not produce the public good and consequently do not pay the corresponding reproductive cost [31–34].

In order to adapt the model to the case of a homogeneous population where all cells produce the public good we absorb the cost of the public good into the baseline division rate. The equation describing the population size is given by (see [23] for details of the derivation):

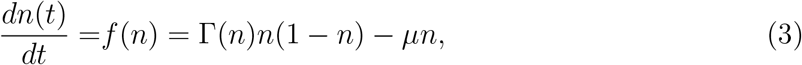

where *n*(*t*) is normalized with respect to the carrying capacity (the maximal population size *N* ^2^) and therefore ranges from 0 to 1, and the division rate Γ(*n*) is density-dependent and given by

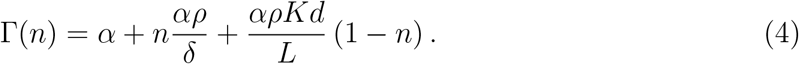

The different terms in the growth rate Γ(*n*) can be given distinct interpretations: the first term is the baseline growth rate in the absence of the GF, the second term is the average GF-contribution from all cells and the last term is an additional growth benefit from the GF due to increased local GF concentration, which is larger for low densities when the factor *n* − 1 is large. This quantity depends on

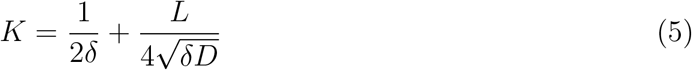

and scales with a factor *d/L* implying that as the cell size *d* decreases (in relation to the system size *L*) the direct benefit is reduced.

### Experimental methods

Cells from the cell line U3065 obtained from the Human Glioma Cell Culture (HGCC) resource [35] were suspended in stem cell medium and plated on 384 well plates (BD Falcon Optilux TC #353962) coated in laminin. Seven different initial densities were used ranging from 2000-30 cells/well and each density was replicated eight times. The cells were cultured at 37 °C and 5 % CO_2_ for 168 hours and imaged using a IncuCyte (modell?) microscope at 20x magnification every second hour. The images were converted to binary and all objects smaller than 100 pixels^2^ were removed. The normalized cell density (degree of confluency) was estimated by calculating the ratio of the total area to the total area of the image. Growth curves for each initial density was calculated by averaging the normalized cell density across all eight replicates. In order to avoid effects of crowding the time series were truncated at 80 hours.

Per-capita growth rates were estimated from the time series by smoothing the curves using a 12 hour moving average and then calculating the numerical derivative

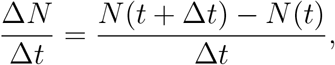

where *N* (*t*) is the number of cells in the smoothed time series and Δ*t* = 2 hours.

The parameters of the ODE-model (3) were estimated by minimising the squared error between the model and the data for all growth curves simulatenously using the error function:

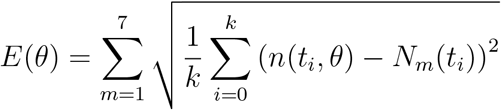

where *k* = 40 is the number of time points *t_i_* and the outer sum runs across the growth curves *N_m_* obtained for different initial densities. The numerical solution of the ODE-model *n*(*t*, *θ*) was obtained by using the normalized density at *t* = 0 as the initial condition and *θ* = (*A*, *B*, *μ*) are the parameters of the model. Numerical solutions were calculated using an Euler-forward scheme with time step of 0.2 hours.

## RESULTS

First, we present results obtained from analyzing an ordinary differential equation (ODE)-model of the system (3). Next, we test the validity of these results by comparing them to outcomes from our individual-based simulation approach. Last, to confirm our theoretical predictions, we analyze experimental results by fitting the ODE-model to time series data of *in vitro* cultured glioblastoma cells.

### Analysis of ODE-model

Depending on the relation between the parameters the ODE-model (3) it can give rise to different dynamics. We will here focus on the impact of the baseline division rate *α* and the death rate *μ*. For *μ* = 0 the system (3) has two non-negative fixed points, *n*^★^ = 0 and *n*^★^ = 1 (see fig. 2A), corresponding to a system void of cells and at carrying capacity respectively. This holds true as long as Γ(*n*) = 0 has no positive solutions, which is the case for the baseline parameters. The presence of a density-dependent division rate leads to non-monotonous per-capita growth rate (*f* (*n*)*/n*) as can be seen in fig. 2B. This is in contrast with the case where Γ(*n*) = constant (i.e. logistic growth) where the per-capita growth rate equals 1 – *n* – *μ* and is a linear and decreasing function of *n*.

**FIG. 2.**
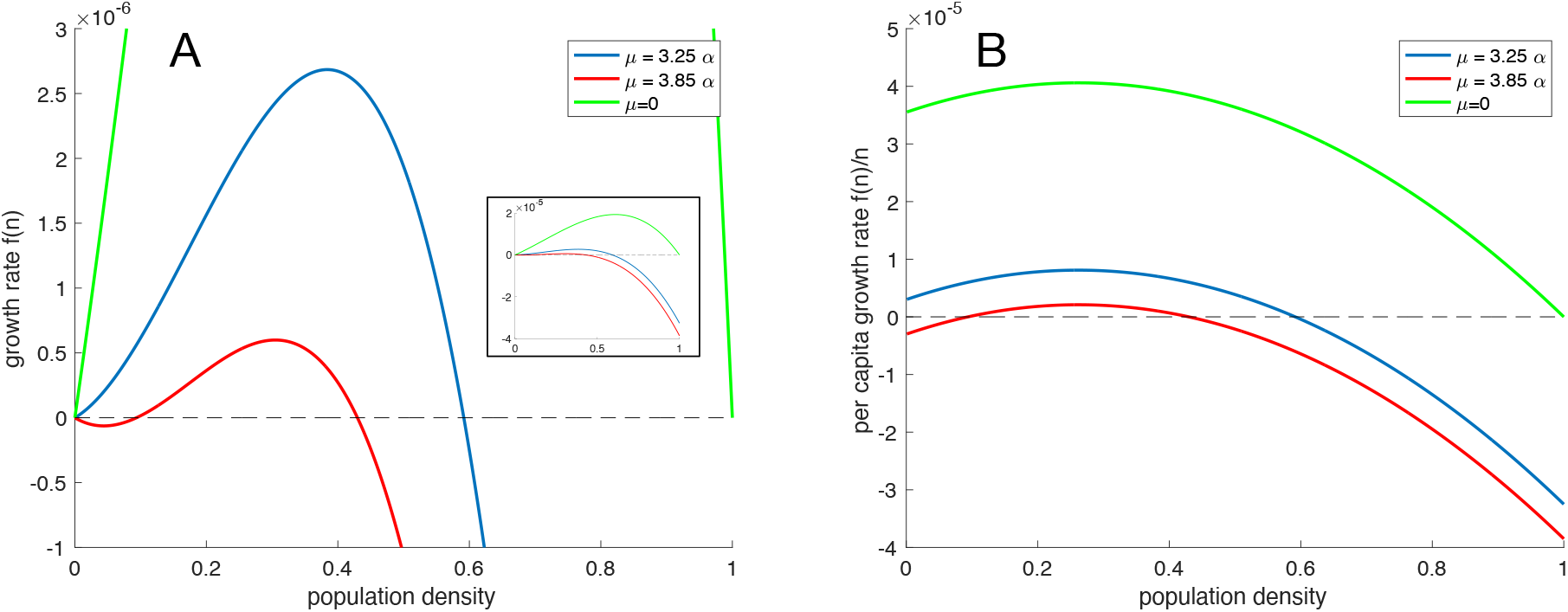
Growth rates. (A) The population growth rate (3) as a function of the population density for three different values of the death rate *μ*. The inset shows the entire range of growth rates. (B) The per-capita growth rate *f* (*n*)*/n* as a function of the population density for three different values of the death rate *μ*. Here the Allee effect is evident as an increasing per-capita growth rate at low densities. All parameter values are given in table I.

For small and vanishing values of *μ*, the per-capita growth rate is an increasing function for small densities, but remains positive. This is known as a weak Allee effect [13], whereas for higher *μ* the per-capita growth rate becomes negative for small densities leading to the extinction of population below a certain critical threshold, which is termed a strong Allee effect [13]. The existence of a strong Allee effect is thus equivalent to the fixed point at the origin *n*^★^ = 0 being stable rather than unstable. The stability can be determined using linear stability analysis and the criterion for stability is that *f*′(0) > 0. We find that the fixed point is stable (or equivalently a strong Allee effect exists) when

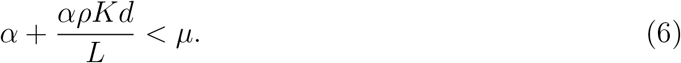

The critical death rate above which we expect to observe a strong Allee effect in the IB-model is thus given by

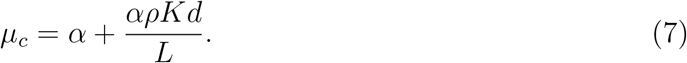

The critical density *n_c_* below which the population is driven to extinction can be calculated explicitly as the unstable interior fixed point of (3), and is given by

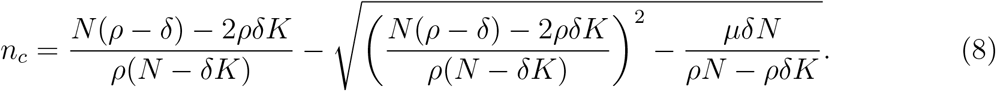

### Comparison to the IB-model

We now move onto comparing our analytical predictions to the IB-model.

#### Long range dispersal

We start by looking at the case when dispersal is long range and newborn cells are dispersed randomly throughout the entire domain. Figure 3A shows a comparison between the IB-model and a numerical solution of the logistic system (3) for three different initial conditions using the baseline parameter values (see table I). Agreement between the IB-model and the ODE is very good and we can conclude that in this scenario autocrine signaling induces a strong Allee effect, since low initial densities give rise to population extinction.

**FIG. 3.**
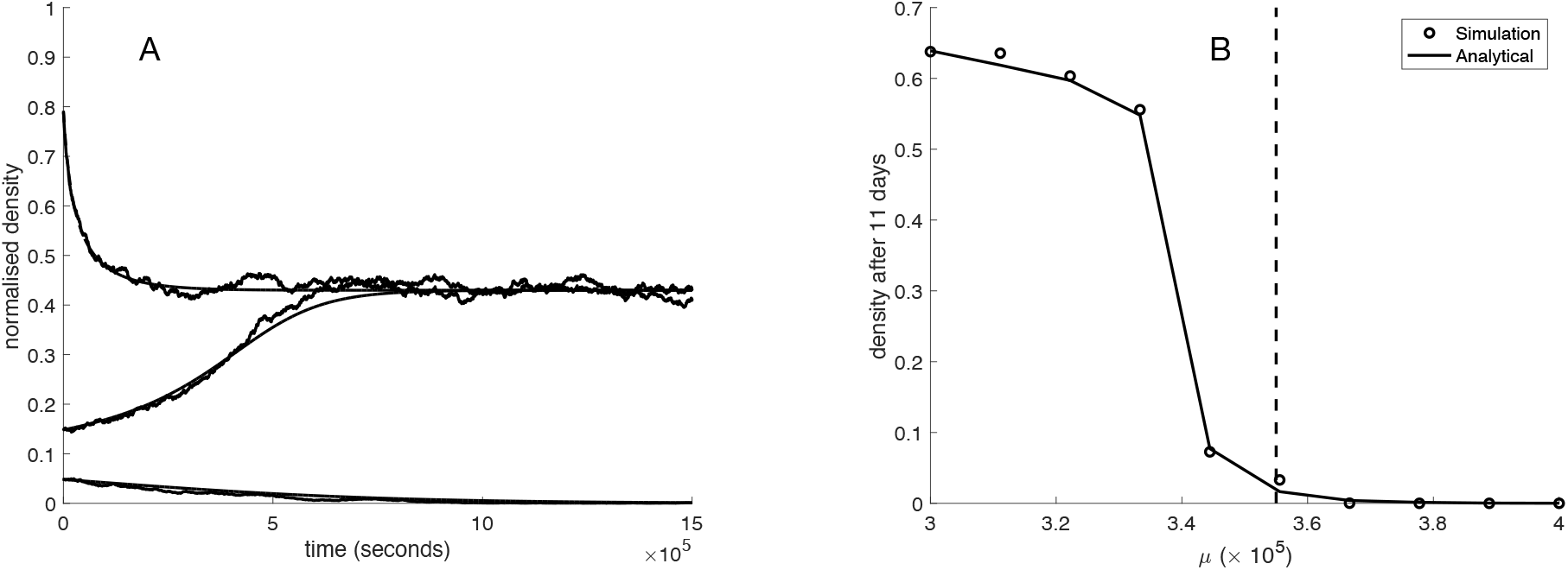
Comparison of the ODE-model and IB-model under long range dispersal. (A) The dynamics of the IB-model (dashed) and the numerical solution of (3) (solid) under long range dispersal for three different initial conditions. All parameter values are given in table I and the death rate *μ* = 3.85 × 10^−5^. (B) The population density after 11 days of the IB-model (circles) and the numerical solution of (3) (solid line) under long range dispersal and no cell migration when the the death rate *μ* is varied. All other parameter values are given in table I.

In order to investigate the impact of the death rate on the Allee effect we initialized the system at a low initial population size of *n*_0_ = 10^−2^ for death rates in the range 2 − 4 × 10^−5^ and ran the IB-model for 11 days and recorded the density of cells. This was compared to the numerical solution of the ODE-model and the result is shown in fig. 3B, where the vertical dashed line corresponds to the theoretically predicted death rate *μ_c_* at which the strong Allee effect emerges. Please note that this value will deviate slightly from the results of the IB-model since the simulations are initialized with small, but non-zero density. A more exact value of the critical death rate can be obtained by setting *n_c_* = *n*_0_ = 10^−2^ in (8) and solving for the death rate *μ*. This expression is however much more complicated than the simple expression for the critical death rate (7).

#### Short range dispersal

We now analyze the case when newborn cells are placed next to the parent cell. Here the migration rate of the cells becomes an important parameter since movement of cells tends to reduce spatial correlations and bring the system closer to the mean-field limit. We therefore start by analyzing the extreme case of short range dispersal and no cell migration, and again compare the long-term dynamics of the IB-model with the numerical solution of the ODE-model (3). By comparing the long-term dynamics we see that the ODE-model severely over-estimates both the population density and the critical death rate *μ_c_* (see fig. 4A). This might seem a bit surprising since local dispersal leads to clumping of cells, and cells in such a configuration would on average experience a higher GF-concentration (compared to an evenly dispersed population) due to the production from neighbouring cells. However, local dispersal also leads to increased competition for space and therefore has a negative effect on the rate of division. This is because cells that are trapped by neighbouring cells cannot divide, which reduces the effective birth rate. This latter effect seems to dominate for the baseline parameter values and as a result the Allee effect becomes even more pronounced.

**FIG. 4.**
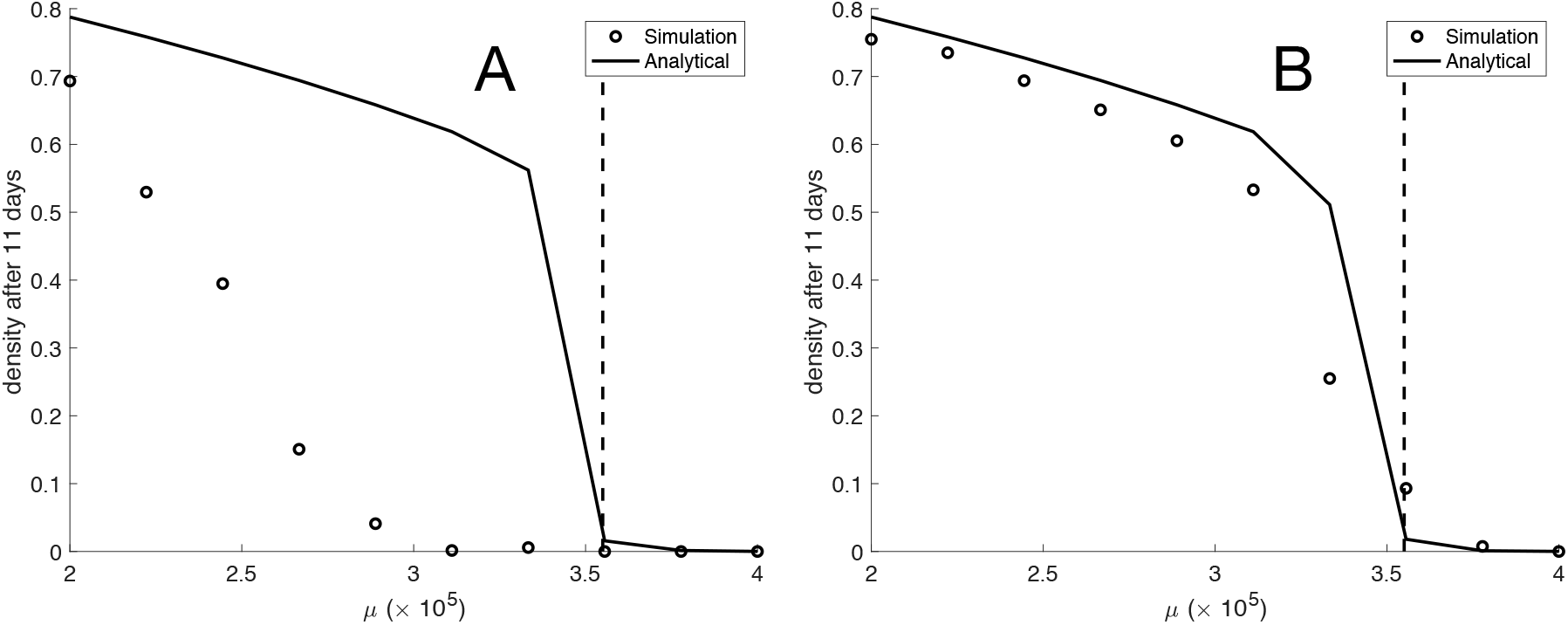
The population density after 11 days as a function of the death rate *μ*. Results from the IB-model (circles) and the numerical solution of (3) (solid line) under (A) short range dispersal and no cell migration and (B) short range dispersal and a migration rate of *ν* = 4 × 10^−4^ s^−1^. All other parameter values are given in table I.

For a realistic migration rate of *ν* = 4 × 10^−4^ s^−1^ corresponding to a cellular diffusion coefficient of 10^−8^ cm^2^/s [30] agreement is considerably better (see fig. 4B). This suggests that experimentally observed migration rates are sufficient to remove the effects of crowding and improve the agreement between the IB-model and the analytical result.

### Comparison to experimental data

Having ascertained that the ODE-model gives an accurate description of the IB-model at realistic rates of cell migration we now move on to fitting the ODE-model to experimental data. We assume a growth rate of the form

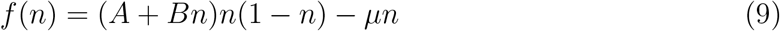

where

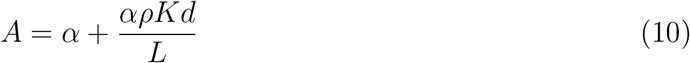

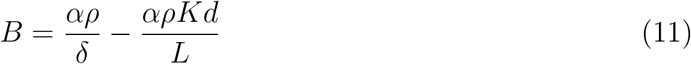

We used least squares minimisation to find numerical values of the constants *A*, *B* and *μ* such that the deviation between the model and growth curves for seven different initial population sizes was minimized. The optimal fit was found for (*A*, *B*, *μ*) = (0.0948, 0.1472, 0.0748) and a comparison between the growth curves and the model dynamics can be seen in fig. 5A. In order to investigate if the data could be explained with a simpler model we also fitted a standard logistic growth function, which corresponds to the special case of zero GF production (*ρ* = 0). The logistic model only has two parameters: a birth rate and a death rate. We found that the logistic model gave larger model error (RMSE of 0.13 compared to 0.11 for the model with Allee effect). In order to account for the larger number of parameters in the ODE-model we also calculated the AIC and found that the ODE-model with an Allee effect has a lower ACI compared to the logistic model (−574 vs. −622). We thus conclude that the ODE-model is a better description of the experimental data compared to the logistic equation.

**FIG. 5.**
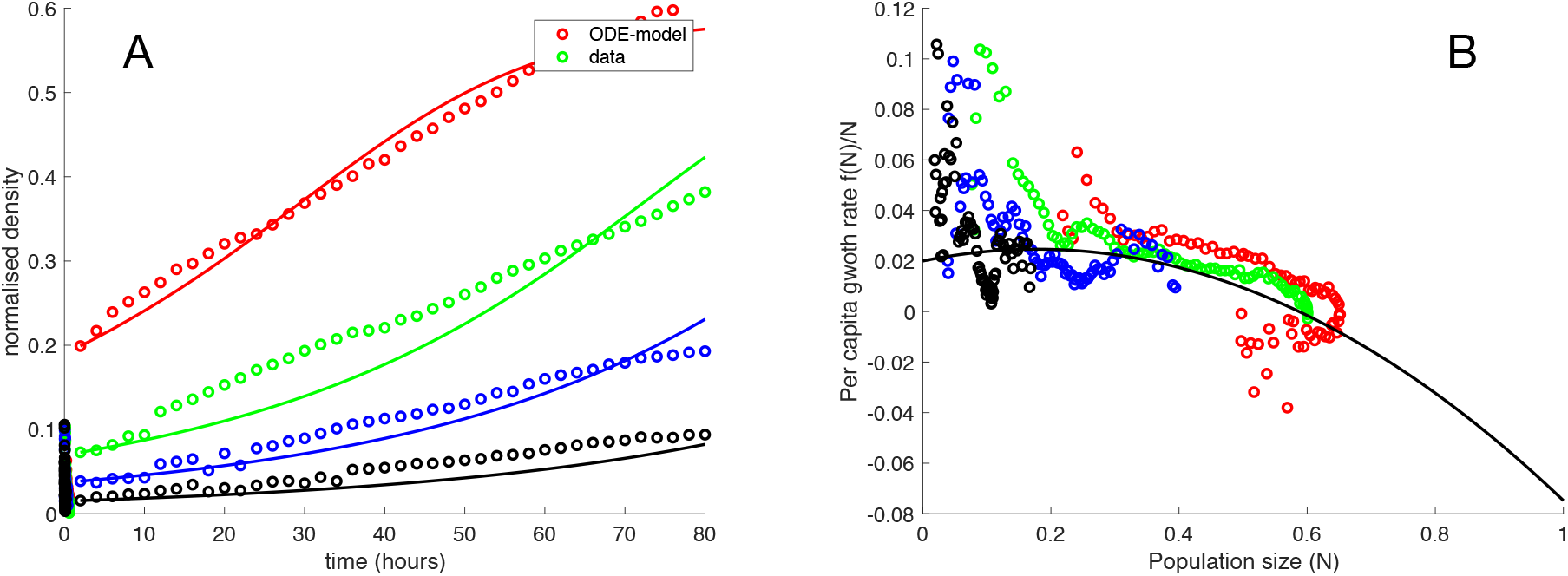
A. Least squares fit of the ODE-model to *in vitro* growth of glioblastoma cells. The optimal fit was found for (*A, B, μ*) = (0.0948, 0.1472, 0.0748). B. per-capita growth rate from experimental data (dots) and the per-capita growth rate from the fitted model (solid line).

## DISCUSSION

We have proposed a mathematical model to explain the Allee effect, a phenomenon observed in many experimental data sets of cancer cell population dynamics. Our model, which posits that paracrine secreted growth factors increase the rate of cell division, yields population dynamics that can exhibit weak or strong Allee effect depending on the relationships between the model parameters. Fitting the model to a dataset of a patient-derived glioma cell line showed that a model with a weak Allee effect provided the best fit.

In recent years many studies have focused on the Allee effect in cancer, either in an effort to understand its origin or to investigate how the effect can be therapeutically exploited [36]. For example, Böttger et al. [37] showed that density-dependent migration rates can give rise to an Allee effect in glioblastoma. Another potential cause of the effect was proposed by Konstorum et al. [38], who investigated the impact of feedback regulation in cancer stem cell dynamics. An explanation in terms of density-dependent proliferation rates was explored by Johnson et al. [18], and similarly by Fadai et al. [39], who both connected the per-capita growth rates in an individual-based model with coefficients in an ODE-model that recapitulated the growth rate decline as the cell populations became smaller. Neither of these results were directly linked to production and consumption of secreted factors among the cell populations.

Here, we provide a mechanistic explanation for a density-dependent proliferation rate in terms of local growth factor concentration. Mathematical analysis of the individual-based model allowed for the derivation of an ODE-model whose coefficients depend on the IBM parameters. Our model was fitted to experimental data, and we found that a weak Allee effect best explained the empirical observations. This is in accordance with the findings of Fadai et al. [39] who also observed a weak Allee effect in breast cancer cell populations. This suggest that a more general pattern of self-interaction-driven negavive feedback among cell lines in which autocrine signaling is present.

In light of cancer therapy, a strong Allee effect would be more beneficial for the ability to control a tumor, since the strong effect leads to population extinction at critically low population densities. Such conditions are typically present after therapeutic interventions (e.g. surgery and radiotherapy). If the population dynamics could be tipped towards a strong Allee effect during therapy, similar to an extinction threshold [40], one could observe decreased risk of recurrence. This critical decline could for example be achieved by reducing the effect of the growth factor by inhibiting its binding to cell surface receptors, or by increasing the factor’s decay rate. Beyond these threshold considerations, the presence of Allee effects has implications for tumor control that needs to consider the emergence of aggressive variants that were previously below detectable thresholds. These variants could receive an advantage, as the previously dominant cancer cell population against which treatment is initially chosen observes fitness decreases due to an Allee effect.

In conclusion, our findings provide a general model for the population dynamics of cancer cells driven by autocrine signaling. We provide a possible mechanistic explanation for the ubiquitous Allee effect. Further, our findings warrant more research into the therapeutic benefits of altering the effects of autocrine signaling to understand and achieve tumor control.

## Notes

### Competing Interest Statement

The authors have declared no competing interest.

## References

[1] A. Laird, British Journal of Cancer (1963).

[2] S. P. Nissley, P. A. Short, M. M. Rechler, J. M. Podskalny, and H. G. Coon, Cell 11, 441 (1977).

[3] G. J. Todaro and J. E. De Larco, Cancer research 38, 4147 (1978).

[4] K. S. Korolev, J. B. Xavier, and J. Gore, Nature Reviews Cancer 14, 371 (2014).

[5] C. C. Maley, A. Aktipis, T. A. Graham, A. Sottoriva, A. M. Boddy, M. Janiszewska, A. S. Silva, M. Gerlinger, Y. Yuan, K. J. Pienta, et al., Nature Reviews Cancer 17, 605 (2017).

[6] B. Bronk, G. Dienes, and R. Johnson, Biophysical journal 10, 487 (1970).

[7] A. Marusyk, M. Janiszewska, and K. Polyak, Cancer cell 37, 471 (2020).

[8] P. Gerlee and P. M. Altrock, Proceedings of the National Academy of Sciences USA 112, E2742 (2015).

[9] P. M. Altrock, L. L. Liu, and F. Michor, Nature Reviews Cancer 15, 730 (2015).

[10] A. Marusyk, D. P. Tabassum, P. M. Altrock, V. Almendro, F. Michor, and K. Polyak, Nature 514, 54 (2014).

[11] A. S. Cleary, T. L. Leonard, S. A. Gestl, and E. J. Gunther, Nature 508, 113 (2014).

[12] W. Allee and E. S. Bowen, Journal of Experimental Zoology 61, 185 (1932).

[13] F. Courchamp, L. Berec, and J. Gascoigne, Allee effects in ecology and conservation (Oxford University Press, 2008).

[14] A. M. Kramer, B. Dennis, A. M. Liebhold, and J. M. Drake, Population Ecology 51, 341 (2009).

[15] H. G. Davis, C. M. Taylor, J. G. Lambrinos, and D. R. Strong, Proceedings of the National Academy of Sciences 101, 13804 (2004).

[16] M. S. Mooring, T. A. Fitzpatrick, T. T. Nishihira, and D. D. Reisig, The Journal of Wildlife Management 68, 519 (2004).

[17] G. M. Luque, T. Giraud, and F. Courchamp, Journal of Animal Ecology 82, 956 (2013).

[18] K. E. Johnson, G. Howard, W. Mo, M. K. Strasser, E. A. B. F. Lima, S. Huang, and A. Brock, 145, 926 (2019).

[19] M. Janiszewska, D. P. Tabassum, Z. Castaño, S. Cristea, K. N. Yamamoto, N. L. Kingston, K. C. Murphy, S. Shu, N. W. Harper, C. G. Del Alcazar, et al., Nature cell biology 21, 879 (2019).

[20] S. Friberg and S. Mattson, Journal of surgical oncology 65, 284 (1997).

[21] Z. Neufeld, W. von Witt, D. Lakatos, J. Wang, B. Hegedus, and A. Czirok, PLoS Computational Biology 13, e1005818 (2017).

[22] I. Nazarenko, S.-M. Hede, X. He, A. Hedrén, J. Thompson, M. S. Lindström, and M. Nistér, Upsala journal of medical sciences 117, 99 (2012).

[23] P. Gerlee and P. M. Altrock, Physical Review E, 1 (2019).

[24] A. R. A. Anderson, Mathematical Medicine and Biology 22 (2005).

[25] E. F. Juarez, R. Lau, S. H. Friedman, A. Ghaffarizadeh, E. Jonckheere, D. B. Agus, S. M. Mumenthaler, and P. Macklin, BMC systems biology 10, 92 (2016).

[26] P. Gerlee and P. M. Altrock, Journal of The Royal Society Interface 14, 20170342 (2017).

[27] L. M. Morimoto, P. A. Newcomb, E. White, J. Bigler, and J. D. Potter, Cancer Epidemiology and Prevention Biomarkers 14, 1394 (2005).

[28] T. Eviatar, H. Kauffman, and A. Maroudas, Arthritis & Rheumatism 48, 410 (2003).

[29] J. V. Nauman, P. G. Campbell, F. Lanni, and J. L. Anderson, Biophysical journal 92, 4444 (2007).

[30] A. R. Anderson and M. Chaplain, Bulletin of Mathematical Biology 60, 857 (1998).

[31] M. Archetti and I. Scheuring, Evolution 65, 1140 (2010).

[32] M. Archetti and I. Scheuring, Journal of Theoretical Biology 299, 9 (2012).

[33] G. J. Kimmel, P. Gerlee, J. S. Brown, and P. M. Altrock, Communications Biology 2, 1 (2019).

[34] B. Johnson, P. M. Altrock, and G. J. Kimmel, Royal Society open science 8, 210182.

[35] Y. Xie, T. Bergström, Y. Jiang, P. Johansson, V. D. Marinescu, N. Lindberg, A. Segerman, G. Wicher, M. Niklasson, S. Baskaran, et al., EBioMedicine 2, 1351 (2015).

[36] M. Delitala and M. Ferraro, AIMS Mathematics 5, 7649 (2020).

[37] K. Böttger, H. Hatzikirou, A. Voss-Böhme, E. A. Cavalcanti-Adam, M. A. Herrero, and A. Deutsch, PLoS Computational Biology 11, e1004366 (2015).

[38] A. Konstorum, T. Hillen, and J. Lowengrub, Bulletin of Mathematical Biology 78, 754 (2016).

[39] N. T. Fadai, S. T. Johnston, and M. J. Simpson, Proceedings of the Royal Society A 476, 20200350 (2020).

[40] G. J. Kimmel, F. L. Locke, and P. M. Altrock, Proceedings of the Royal Society B 288, 20210229 (2021).

